# Shape transformation of vesicles induced by orientational arrangement of membrane proteins

**DOI:** 10.1101/2024.09.28.615559

**Authors:** Menglong Feng, Kunhao Dong, Yuansheng Cao, Rui Ma

**Author notes:** These authors contributed equally to this work (Menglong Feng, Kunhao Dong).

## Abstract

Vesicles of lipid bilayer can adopt a variety of shapes due to different coating proteins. The ability of proteins to reshape membrane is typically characterized by inducing spontaneous curvature of the membrane at the coated area. BAR family proteins are known to have a crescent shape and can induce membrane curvature along its concaved body axis but not in the perpendicular direction. We model this type of proteins as a rod-shaped molecule with an orientation and induce normal curvature along its orientation in the tangential plane of the membrane surface. We show how a ring of these proteins reshape an axisymmetric vesicle when the protein curvature or orientation is varied. A discontinuous shape transformation from a protrusion shape without a neck to a one with a neck is found. Increasing the rigidity of the protein ring is able to smooth out the transition. Furthermore, we show that varying the protein orientation is able to induce an hourglass-shaped neck, which is significantly narrower than the reciprocal of the protein curvature. Our results offer a new angle to rationalize the helical structure formed by many proteins that carry out membrane fission functions.

## I. INTRODUCTION

Cells are able to remodel their shapes to perform specific biological functions. A small vesicle buds off from the plasma membrane during endocytosis [1–3]. A mother cell is divided into two daughter cells during mitosis [4, 5]. A crawling cell forms protrusions at the leading edge when migrating on a substrate [6, 7]. The various morphology of cells is attributed to proteins that either induce membrane curvatures or produce forces to deform the membrane [8–15]. Clathrin molecules can assemble into a cage-like structure and scaffold the membrane into a spherical vesicle [16, 17]. BAR family proteins, initially identified in yeast cells [18, 19], regulate the shape of membranes in various biological processes, including endocytosis [14, 20, 21], autophagy [22], cell division [23], motor transportation [24] and membrane re-modelling [25]. Meanwhile, their interaction with other cellular structures such as plasma membrane [26], cytoskeletons [23, 27, 28], and dynamin [6, 29] can play an important role [30–32]. Due to the crescent shape of the BAR proteins, they tend to curve the membrane along the direction of their body axis, thus remodelling the membrane into a tubular shape [33, 34]. Even though different BAR proteins vary in their length, intrinsic curvature, and binding affinity to the membrane [35, 36], the ability to induce membrane tubulation is a common property for most of the BAR proteins [37–41]. The classical BAR and N-BAR proteins containing a unique N-terminal amphipathic helix [42, 43] can generate tubules, with diameters from 20 to 60 nm [44], and bend the membrane towards the leaflet that they are embedded. F-BAR proteins can induce wider tubules from 60 to 100 nm [45, 46] in diameter and also bend the membrane towards the attached leaflet. By contrast, I-BAR proteins induce curvatures towards the unattached leaflet [47, 48]. When bound to the inner leaflet of a closed vesicle, I-BAR proteins are able to produce membrane protrusions [49]. The ability of BAR proteins to remodel the membrane depends on the protein density [50–53], as well as the mechanical properties of the membrane [54], such as membrane tension and bending rigidity [55–57]. BAR proteins can scaffold membranes into tubular shapes at high density. However, at low density, they bind to sites with preferred curvatures on the membrane, acting as a curvature sensor [58–63].

The organization of BAR proteins also plays an important role when generating membrane curvatures [64, 65]. Computer simulations suggest that BAR proteins form a helical string-like aggregate to facilitate membrane fission. As a curved rod-shaped molecule [66], the orientation of the BAR proteins along the rod axis is tilted relative to the circumferential direction of the tubular membrane. Whether the titled angle in the helical string of the protein assembly plays a role in generating membrane curvatures remains unclear.

In this paper, we study the deformation of a spherical vesicle induced by proteins that produce normal curvature along a tangential direction on the membrane and focus on how the orientation of the proteins influence the shape of the vesicle. The paper is organized as follows. Firstly, we will illustrate the geometry of an axisymmetric surface and formulate our theoretical model of lipid membrane coated by anisotropic proteins. Secondly, we will show the membrane shapes acquired by our numerical method, where shape diagrams with respect to the preferred curvature and protein orientation are displayed to reveal the shape transformation. Thirdly, we will elaborate how other factors including coating position and the bending rigidity influence the cellular morphology and shape transformation. Finally, we will summarize this work and compare our results with similar research on anisotropic proteins.

## II. MECHANICAL MODEL OF MEMBRANE AND PROTEINS

In this section, we briefly introduce our theoretical model. For mathematical details, one can refer to the Appendix. We consider a closed membrane with axisymmetric shape, whose cross section is parameterized with [*r*(*u*), *z*(*u*)] (Fig. 1). A dimensionless coordinate *u* ∈ [0, 1] is introduced to label the meridional position on the membrane with *u* = 0 and *u* = 1 indicating the north pole and the south pole respectively. With the help of two auxiliary variables *ψ*(*u*) and *h*(*u*) that satisfy the geometric relations in Eqs. (A2) and (A3) (see Appendix A), the two principal curvatures of the membrane surface can be simplified as

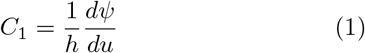

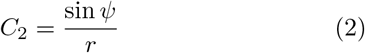

**FIG. 1.**
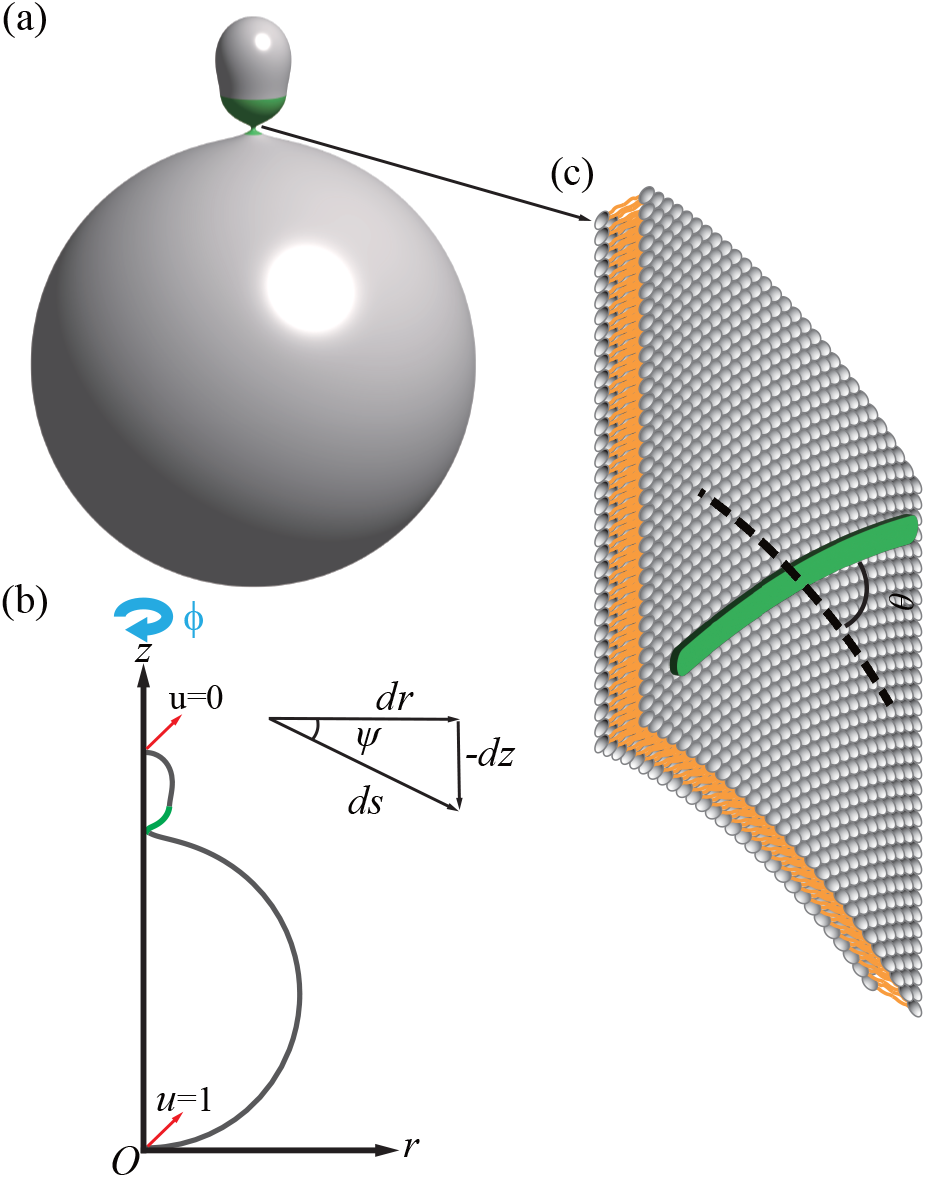
(a) Schematic illustration of the neck formed by coating proteins that are tilted relative to the azimuthal direction on the membrane. The proteins are assumed to be rod-shaped and tend to bend the membrane along their body orientation. (b) Geometric representation of a closed axisymmetric surface. The green part represents the protein coated region which forms a ring.

We assume a closed membrane with a conserved area 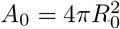 and employ Helfrich theory to model the free energy of the membrane as

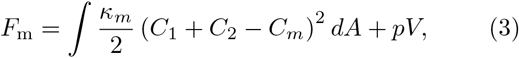

where *κ*_*m*_ denotes the bending rigidity of the membrane, *C*_*m*_ denotes the isotropic spontaneous curvature [67]. The pressure difference between the inside and outside of the vesicle is denoted by *p*. We fix the pressure 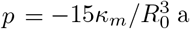 and allow the volume of the vesicle to change. The negative sign implies that the inner pressure is higher than the outer pressure. Considering the typical value for the membrane bending rigidity *κ*_*m*_ = 20 *k*_*B*_*T* and the size of a vesicle of 100 nm in diameter, the pressure difference *p* is about 0.01 MPa.

The rod-shaped BAR proteins are assumed to lay on the membrane surface and orient relative to the circumferential direction with a titled angle *θ* (Fig. 1). From Euler’s formula [68], the normal curvature along the *θ* direction reads

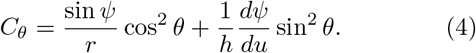

Deviation of the normal curvature *C*_*θ*_ from the proteins’ preferred curvature *C*_*p*_ leads to the anisotropic bending energy of the BAR proteins

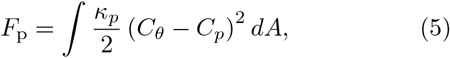

where *κ*_*p*_ denotes the bending rigidity of the protein coat. To label material points on the membrane surface, we introduce the function *A*(*u*), representing the surface area covered from the north pole (*u* = 0) to the circumference with coordinate *u*. We assume the protein only covers a tiny fraction (1%) of the membrane area and form a ring at the region *A*_1_ *< A*(*u*) *< A*_2_. The coating area of proteins is *A*_coat_ = *A*_2_ − *A*_1_, and the coating position is indicated by the average (*A*_1_ + *A*_2_)*/*2.

Summing up the contributions from both the membrane *F*_m_ and the proteins *F*_p_, as well as Lagrangian multipliers *F*_L_ to enforce geometric relations between the shape variables, we express the total free energy of the system as

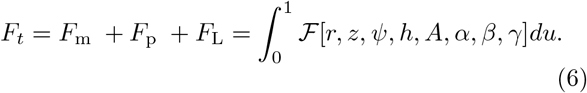

Note that *F*_*t*_ is a functional of the geometric variables *r, z, ψ, h, A*, and the Lagrangian multipliers *α, β, γ* (see Appendix A for the detailed expression).

In particular, the Lagrangian multiplier *γ* serves as the membrane tension to enforce *dA/du* = 2*πrh*. We then employ the variational method to obtain the Euler-Lagrange equations and the associated boundary conditions, which are numerically solved with the MATLAB solver bvp5c that is designed for boundary value problems defined by ordinary differential equations.

## III. RESULTS

### A. Shape transformations by a protein ring near the north pole

We first study the shape transformations of the spherical vesicle when the protein ring is located near the north pole with (*A*_1_ + *A*_2_)*/*(2*A*_0_) = 0.033. Because a soft protein coat cannot dramatically deform the spherical vesicle (data not shown), we set the bending rigidity of the protein ring twice as rigid as the membrane, i.e., *κ*_*p*_ = 2*κ*_*m*_. When the the titled angle is relatively high, for instance, *θ* = 46°, continuously varying the protein curvature *C*_*p*_ transforms the spherical vesicle into a vase-like shape with the protein coated region grow outward and form a high positive curvature bump (Fig. 2a). This is because the proteins tend to produce positive curvature along the meridional direction. However, with a relatively small tilted angle *θ* = 22°, two types of shapes emerge as *C*_*p*_ goes up. A protrusion, which is characterized with a region with negative curvature *C*_1_, is formed at relatively small *C*_*p*_ (Fig. 2b, shapes labeled with A, B, C). Another type of solution is featured with a shape in which the protein ring is contracted inward to form a protrusion with a narrow neck (Fig. 2b, shapes labeled with D, E, F). The two types of shapes are connected with a Gibbs triangle in the total free energy curve, which indicates a first-order (discontinuous) phase transition at the crossing point (Fig. 2, pink curve, *C*_*p*_*R*_0_ = 18.81).Note that on the branch from shape *D* to *F* in Fig. 2b, the neck induced by the proteins grows narrower and longer with increasing *C*_*p*_. The shape resembles the tubulation induced by N-BAR proteins observed in experiments [69]. This is because proteins are aligned almost in parallel with the azimuthal direction and tend to contract the membrane to a radius that is close to their preferred curvature *C*_*p*_.

**FIG. 2.**
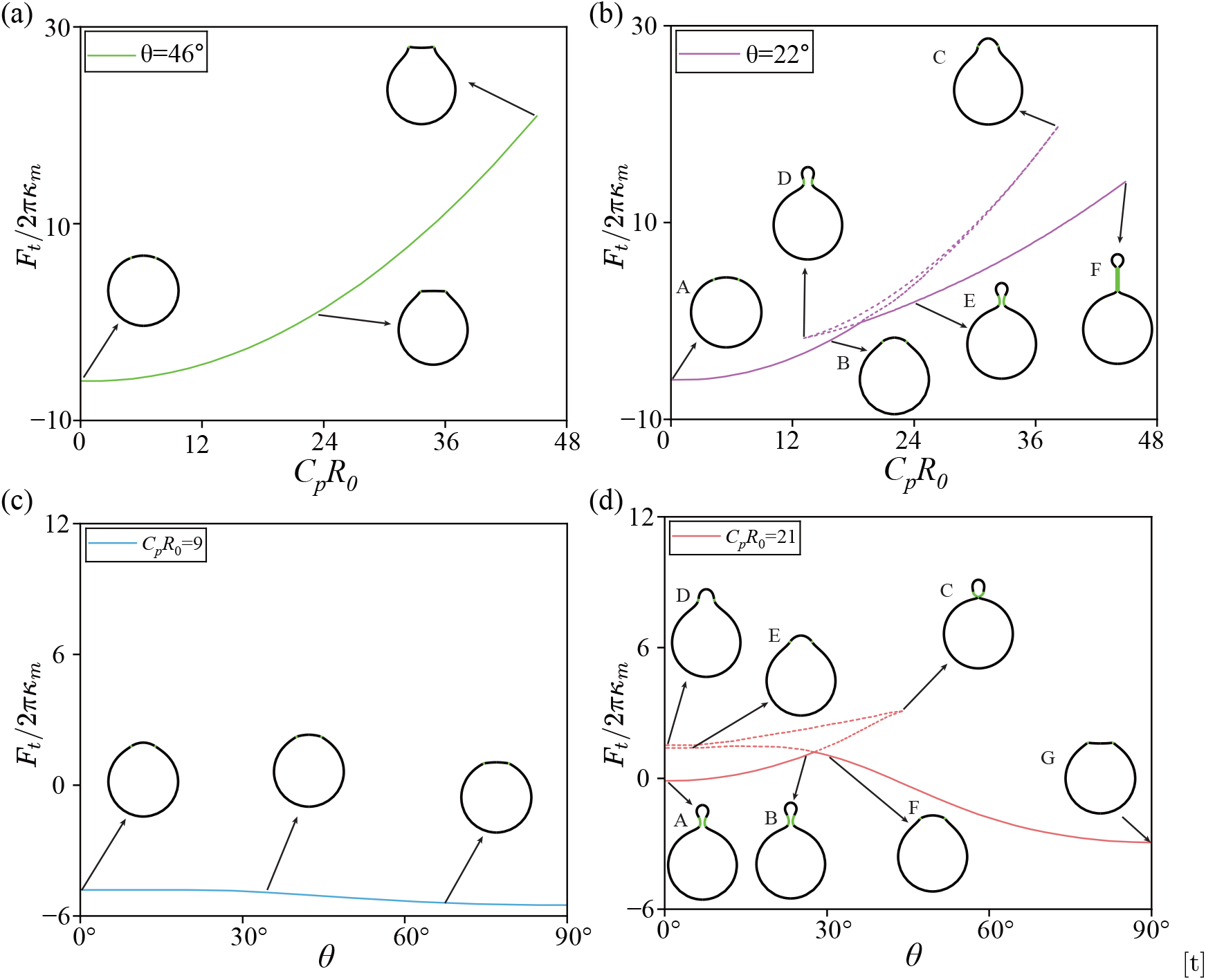
Free energy curves and the associated shape transformations of a vesicle induced by varying the protein curvature *C*_*p*_ in (a, b) or the protein orientation *θ* in (c, d). The solid lines indicate the minimum free energy state and the dash lines indicate states with higher free energy. The protein ring is located near the north pole where (*A*_1_ + *A*_2_)*/*(2*A*_0_) = 0.033, and has bending rigidity *κ*_*p*_*/κ*_*m*_ = 2.

We now investigate how continuously varying the titled angle *θ* transform the membrane shapes. When the protein curvature *C*_*p*_ is small, e.g., *C*_*p*_*R*_0_ = 9, increasing *θ* from 0° to 90° only slightly changes the vesicle shape such that the protrusion formed at small *θ* is flattened or even bent inward at large *θ* (Fig. 2c). If the protein curvature *C*_*p*_ is raised to *C*_*p*_*R*_0_ = 21, at relatively small *θ*, a single *θ* corresponds to three solutions of protrusion shapes, two of which have no neck and the third one has a neck. It is intriguing to see that as *θ* goes up, the neck is narrowed down to a very small radius (Fig. 2d, see shapes labeled with A, B and C), which could induce a small vesicle budding off the membrane. This necking phenomena will be further discussed later in the paper.

Meanwhile, the branch without necks goes from a protrusion shape to a vase-like shape such that the bulge is flattened (Fig. 2d, see shapes labeled with E, F and G). The two branches cross at a critical point in a Gibbs triangle, which indicates a first-order phase transition from a protrusion with neck to a protrusion without neck with increasing *θ* (Fig. 2d, shapes labeled from B to F).

To systematically understand the shape transformations induced by the protein curvature *C*_*p*_ and the tilted angle *θ*, we show the phase diagram in Fig. 3. The phase space is divided into two regions, depending on whether the protrusions have a neck or not. Crossing the boundary from one region to the other represents a first-order phase transition characterized by the Gibbs triangle exemplified in Fig. 2b and d. As *θ* goes from 0° (parallel to the azimuthal direction) to about 45°, an increasing protein curvature *C*_*p*_ is needed for necking to occur. Within the necking region, the radius of the neck is reduced with increasing *C*_*p*_, while the length of the neck is elongated (Fig. 3, dots in purple). Beyond *θ* = 45°, necking cannot occur any more.

**FIG. 3.**
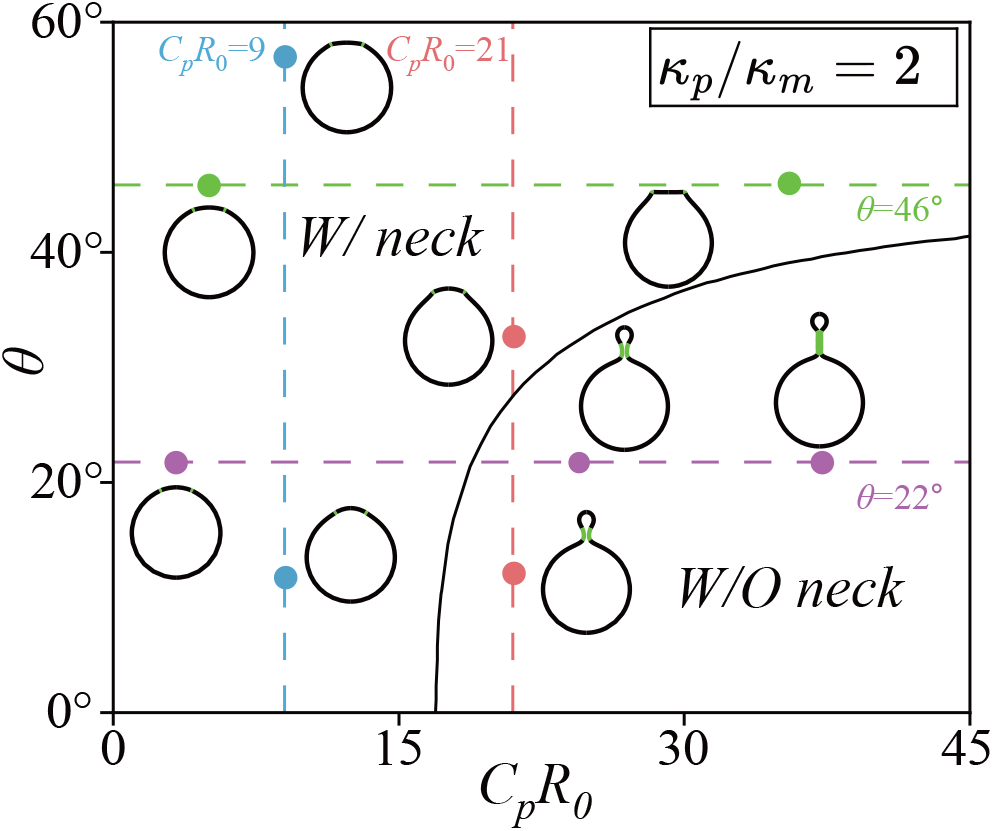
Phase diagram of vesicle shapes as a function of the protein curvature *C*_*p*_ and the protein orientation *θ*. The solid line represents the boundary that separates the region of a protrusion shape without a neck from a region with a neck. Crossing the boundary leads to a discontinuous transition of the vesicle morphology. The four colored dashed lines represent the situation studied in Fig. 2. The protein ring is located near the north pole where (*A*_1_ + *A*_2_)*/*(2*A*_0_) = 0.033, and has bending rigidity *κ*_*p*_*/κ*_*m*_ = 2.

### B. Shape transformations by a protein ring at the equator

We now place the protein ring at the equator, i.e., (*A*_1_ + *A*_2_)*/*(2*A*_0_) = 0.5, and investigate the deformation of the spherical vesicles when varying *C*_*p*_ and *θ*. If *θ* takes a value that is larger than 45°, e.g., *θ* = 46°, varying *C*_*p*_ only slightly alters the spherical vesicle into a rugby shape (Fig. 4a). However, when *θ* is small, such that the proteins are oriented mainly in parallel with the azimuthal direction, we observe two types of solutions connected by a Gibbs triangle in the free energy curve (Fig. 4b). On one of them (Fig. 4b, shapes labeled with A, B and C), shape transformations are similar to that of *θ* = 46°. While on the other (Fig. 4b, shapes labeled with D, E and F), the spherical vesicles are strongly contracted in the equator, resembling a cell division. If the protein curvature *C*_*p*_ is fixed at a small value, e.g., *C*_*p*_*R*_0_ = 9, gradually increasing *θ* almost does not change the vesicle shape (Fig. 4c). However, for large enough *C*_*p*_, a division shape minimizes the total free energy at smaller *θ* and a rugby shape becomes the energy minima at larger *θ* (Fig. 4d).

**FIG. 4.**
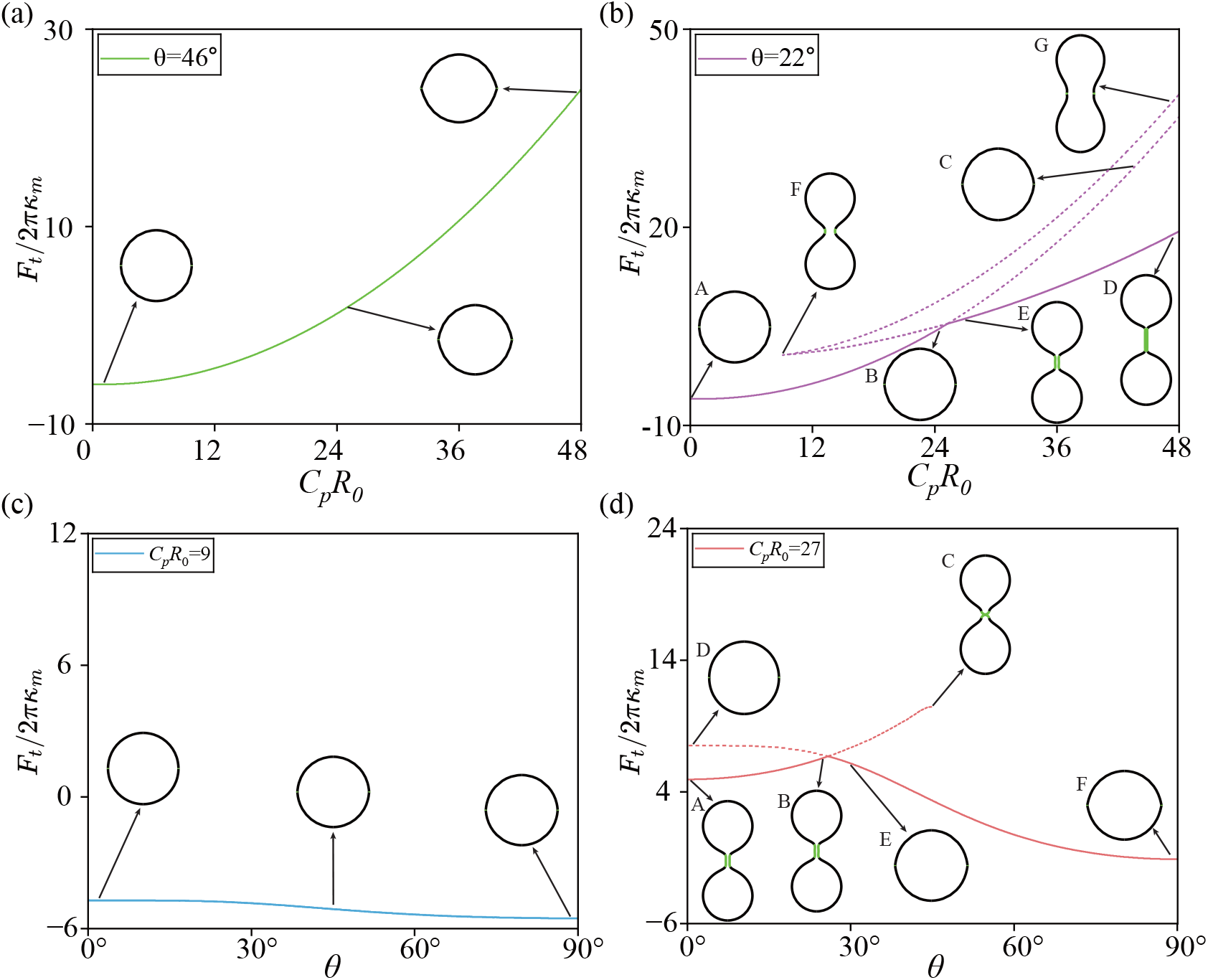
Free energy curves and the associated shape transformations of a vesicle induced by varying the protein curvature *C*_*p*_ in (a, b) or the protein orientation *θ* in (c, d). The solid lines indicate the minimum free energy state and the dash lines indicate states with higher free energy. The protein ring is located at the equator where (*A*_1_ + *A*_2_)*/*(2*A*_0_) = 0.5, and has bending rigidity *κ*_*p*_*/κ*_*m*_ = 2.

The shape transformation behaviors are summarized in the phase diagram in which the division shapes are separated from the rugby shapes by a boundary curve. Crossing the boundary will induce a first order transition of the vesicle shapes. Compared with the case when the protein ring is near the north pole, the boundary curve is shifted towards larger *C*_*p*_ when the protein ring is located at the equator (compare the boundary in Fig. 3 and Fig. 5).

**FIG. 5.**
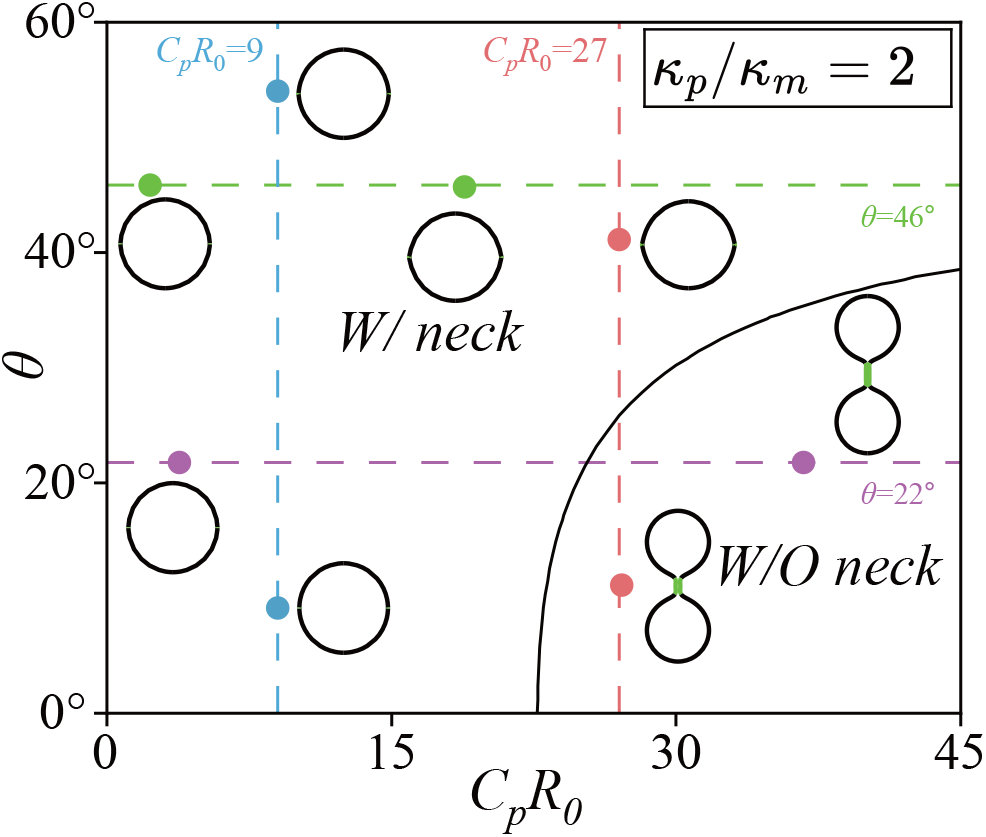
Phase diagram of vesicle shapes as a function of the protein curvature *C*_*p*_ and the protein orientation *θ*. The solid line represents the boundary that separates the region of a protrusion shape without a neck from a region with a neck. Crossing the boundary leads to a discontinuous transition of the vesicle morphology. The four colored dashed lines represent the situation studied in Fig. 4. The protein ring is located at the equator where (*A*_1_ + *A*_2_)*/*(2*A*_0_) = 0.5, and has bending rigidity *κ*_*p*_*/κ*_*m*_ = 2.

### C. Rigid protein coats smooth out the shape transformation

In this section, we fix the position of the protein ring near the north pole and the orientation of proteins at *θ* = 22°, and show the effects of protein rigidity *κ*_*p*_ on shape transformations of the vesicle. When *κ*_*p*_*/κ*_*m*_ = 2, as we have shown in Fig. 2b, with an increasing protein curvature *C*_*p*_, the free energy curve goes through a Gibbs triangle, which indicates a discontinuous shape transition at the crossing point from a protrusion without a neck to that with a neck (see Fig. 6a). With a more rigid protein coat *κ*_*p*_*/κ*_*m*_ = 10, the size of the Gibbs triangle in the free energy curve is reduced and the transition point is shifted towards smaller curvature values (compare Fig. 6a and b). Upon further increasing the protein rigidity to *κ*_*p*_*/κ*_*m*_ = 79, the Gibbs triangle disappears and the shape transformation becomes continuous (see Fig. 6c). The transition point is further shifted towards a smaller protein curvature *C*_*p*_.

**FIG. 6.**
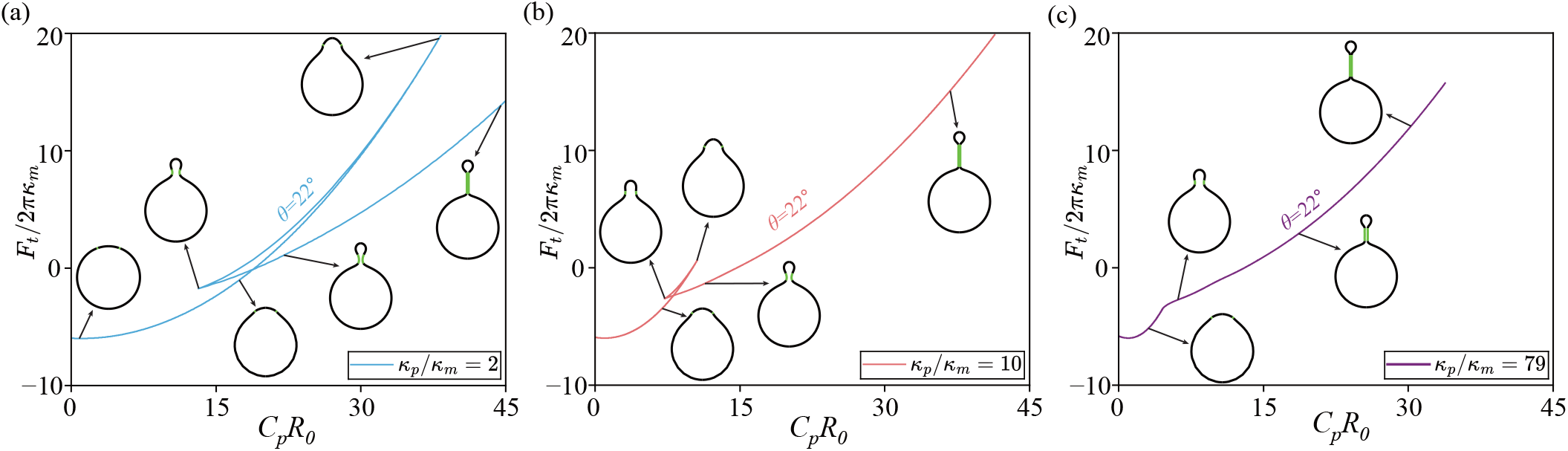
Free energy curves and the associated shape transformations of a vesicle induced by varying the protein curvature *C*_*p*_ for different protein rigidity. From (a) to (c), the protein rigidity *κ*_*p*_*/κ*_*m*_ = 2, 10, 79, respectively. The protein ring is located near the north pole where (*A*_1_ + *A*_2_)*/*(2*A*_0_) = 0.033. The protein orientation is fixed at *θ* = 22°.

### D. Rotating the protein orientation significantly reduces the neck radius

We have shown that upon increasing the protein curvature *C*_*p*_, a smaller vesicle can grow out from a larger vesicle and the neck that connects the two vesicles grows narrower and longer at the same time (see the green part on the vesicle in Fig. 2b). We have also shown that varying the protein orientation *θ* is able to reduce the neck radius (Fig. 2d, from A to B to C). In Fig. 7a, we plot the neck radius as a function of the tilted angle *θ* (Fig. 7a, blue curve). A sharp drop of the neck radius is observed when *θ* approaches about 45°. Compared with the long tube-shaped neck formed by increasing *C*_*p*_, the neck formed by varying the protein orientation *θ* is much shorter and exhibits an hourglass shape, which is similar to previous work on BAR proteins [70]. More strikingly, the neck radius is reduced to a value that is smaller than the reciprocal of the protein’s preferred curvature (indicated by the blue and red dots on the y-axis in Fig. 7). Similar necking morphology difference is also observed when the vesicle exhibits a division-like shape by a protein ring located at the equator (Fig. 7a, red curve). Compared with protein orientation-induced necking, the constriction of the neck radius induced by protein curvature is more gentle and the neck shows a tubular shape stays above the reciprocal of the protein curvature (Fig. 7b). Note that when the tilted angle *θ* is closed to 45°, the necking solution has higher free energy than the one without a neck (plotted in dashed line in Fig. 7). The discontinuous shape transformation occurs at a smaller angle than 45°. However, this does not exclude the physical existence of the necking solution.

**FIG. 7.**
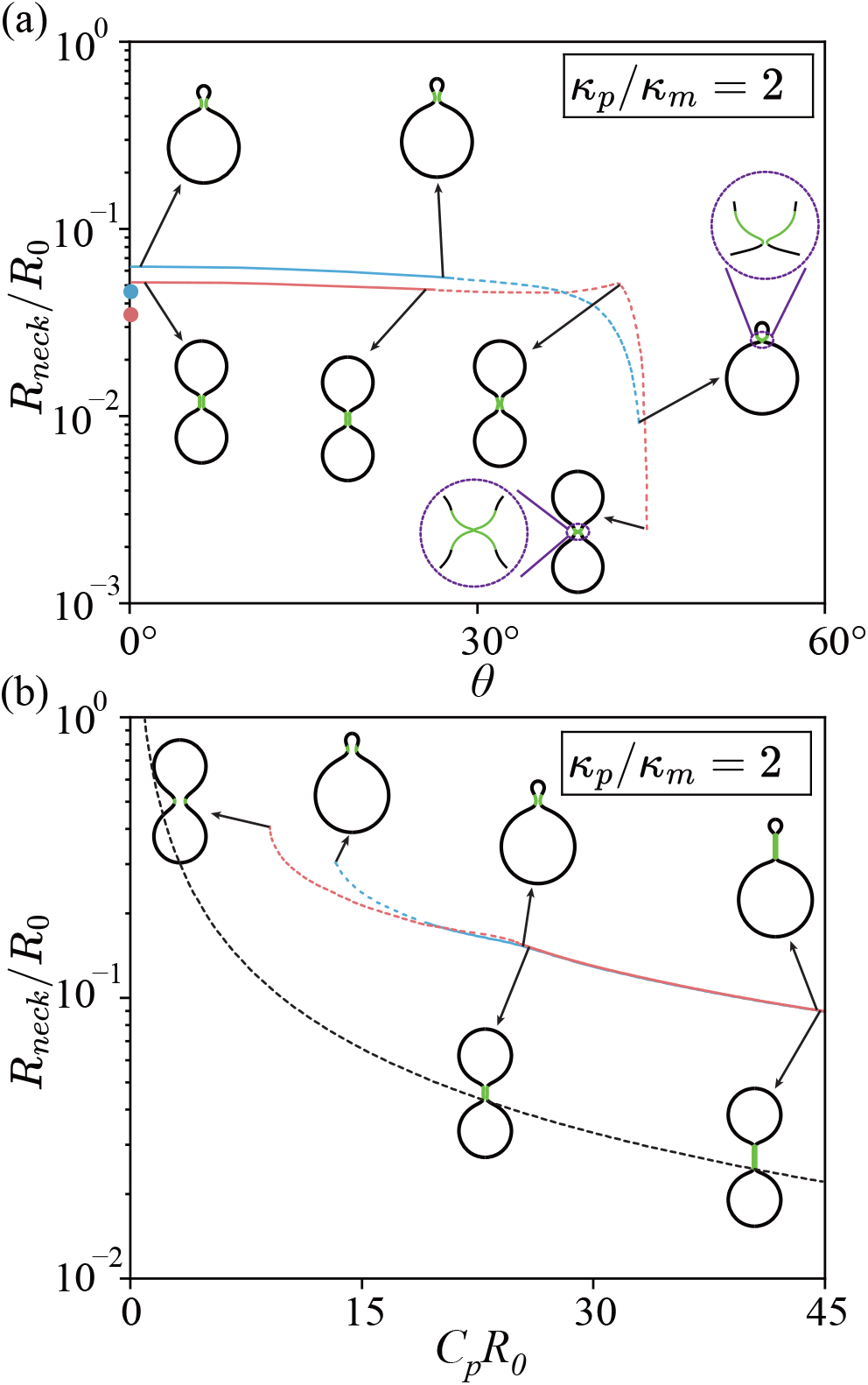
Contraction of the neck radius induced by varying the protein orientation in (a) and the protein curvature in (b). The blue curve represents the situation that the protein ring is located near the north pole where (*A*_1_ + *A*_2_)*/*(2*A*_0_) = 0.033. The red curve represents the case that the protein ring is located at the equator where (*A*_1_ + *A*_2_)*/*(2*A*_0_) = 0.5. The solid part indicate the free energy minimum state. The dotted part indicate metastable states which are locally stable but globally unstable. The blue dot and the red dot on the y-axis in (a) indicate the position of the reciprocal of the protein curvature 1*/*(*C*_*p*_*R*_0_) used to calculating the blue curve and the red curve, respectively. The black dotted curve represent the reciprocal of the protein curvature 1*/*(*C*_*p*_*R*_0_).

In fact, the necking solution represents a metastable state. If *θ* starts from a smaller value of *θ* and is gradually increased to 45°, the vesicle shape can remain on the necking branch provided transition to the other branch needs to overcome a large energy barrier, known as hysteresis effect.

## IV. DISCUSSION

Necking phenomena are common in cellular processes such as endocytosis and cell division. A vesicle is budding off from the plasma membrane when endocytosis occurs [26]. In mammallian cells, the budding is generally attributed to the dynamin protein that is observed to form a helical band around the neck [10, 71]. The chemical energy released by GTP hydrolysis is used to power the constriction of the dynamin. In yeast cells, the role of BAR proteins in driving membrane fission remains controversial [31, 37, 72, 73]. There are studies suggesting BAR proteins can actively constrict the neck of the endocytic pit, therefore driving membrane fission [74, 75]. It is also reported that BAR proteins recruit dynamin to the neck of the membrane buds to facilitate membrane scission [76]. Our study provides a new angle from the protein orientation to explain the constriction of membrane neck by dynamin or BAR proteins. The fact that they are organized into a helical string, in which the proteins are tilted relative to the azimuthal direction, has a physical reason behind it. The preferred radii of BAR family proteins is typically greater than 10nm. However, in order to pinch off the membrane neck, the neck radius needs to be reduced to ∼ 5nm, the same as the thickness of lipid bilayer. Our results suggest that only in a titled organization can the protein ring constrict the membrane neck to a value that is smaller than the protein curvature. Therefore, the constriction function of dynamin or BAR proteins powered by GTP hydrolysis is probably due to reorganization of the helical structure into a more tilted manner.

In this paper, we study how the protein orientation influences the shape of a vesicle when the proteins are organized into a ring on the vesicle. We show that when the protein coat is relatively rigid, varying the protein curvature or the protein orientation induces a discontinuous shape transformation of the vesicle from a protrusion without a neck to that with a neck. A very rigid protein coat can smooth out the transition and move the transition point towards a smaller protein curvature. Furthermore, we show that varying the protein orientation alone is able to induce an hourglass-shaped narrow neck. Our study provides a rationale of the helical band formed by BAR proteins or dynamin proteins when they carry out membrane fission functions during endocytosis. Our study also reveals the importance of the organization of the protein aggregates in reshaping membrane, which is typically neglected in previous researches.

## V. ACKNOWLEDGMMENTS

We acknowledge financial support from the Fundamental Research Funds for Central Universities of China under Grant No. 20720240144 (RM) and the National Natural Science Foundation of China under Grants No. 12374213 (YC) and 111 Project No. B16029.

## Appendix A Description of the geometry of the surface

Here we provide a detailed description of the geometric representation of the vesicle shape, which is parameterized as

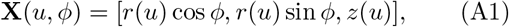

where *ϕ* ∈ [0, 2*π*] is the azimuthal coordinates, *u* ∈ [0, 1] is the meridional coordinates, *r*(*u*) and *z*(*u*) describe the shape of the axisymmetric surface. In order to simplify the expression for the two principal curvature, we introduce two auxiliary functions *ψ*(*u*) and *h*(*u*), which fullfil the geometric relations

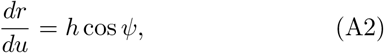

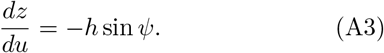

With these notations, the two principal curvatures *C*_1_ and *C*_2_ can be expressed with Eqs.(1) and (2). The function *ψ*(*u*) represents the angle spanned between the tangential direction along the meridian and the radial direction. As for the function *h*(*u*), we require *dh/du* = 0 which means *h*(*u*) is a constant. Because of the relation 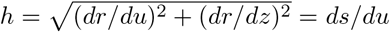, where *s*(*u*) denotes the arclength calculated from the north pole *u* = 0, *h* is equal to the total arclength *S*_*t*_ from the north pole to the south pole. Besides, we also introduce the auxiliary functions *A*(*u*) to denote the surface area ranging from the north pole to a circumferential line with the coordinate *u*. It satisfies the differential equation

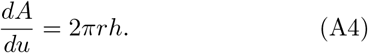

A conserved total surface area *A*_0_ can be easily expressed as the boundary condition for *A*(*u*) with *A*(0) = 0 and *A*(1) = *A*_0_.

The geometric relations expressed by Eqs. (A2), (A3) and (A4) are imposed by introducing three Lagrangian multiplier functions *α*(*u*), *β*(*u*) and *γ*(*u*) in the functional,

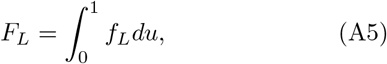

in which

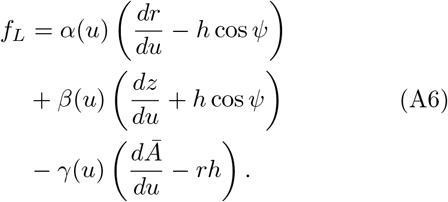

Here *Ā* = *A/*(2*π*) is the reduced surface area function. With these Lagrangian multipliers, the geometric variables *r*(*u*), *z*(*u*), *ψ*(*u*), *h*(*u*) and *A*(*u*) become independent. Note that *γ*(*u*) serves as the membrane tension. If the membrane is inhomogeneous with *A*(*u*)-dependent membrane properties, variation against *A*(*u*) leads to an equation about *dγ/du*. The same equation is also derived in [77].

## Appendix B Helfrich model of the membrane

Eq. (3) describes our model of the membrane. It consists of the Helfrich bending energy and the pressure energy. In the Helfrich bending energy, we neglect the contribution from the Gaussian curvature term, 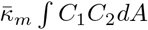, which is a constant for a vesicle of spherical topology due to the Gauss-Bonnet theorem [68]. As for the pressure energy, we allow the volume *V* of the vesicle to change, and fix the parameter *p* which serves as the pressure difference between the outside and inside of the vesicle (*p* = *p*_out_ *p*_in_). In summary, the explicit expression for the Helfrich bending energy reads

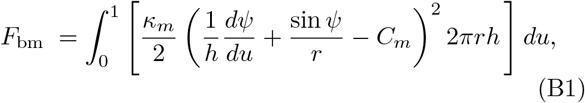

where the bending rigidity *κ*_*m*_ is assume to be a constant and the spontaneous curvature *C*_*m*_ is set to be zero. The pressure energy reads

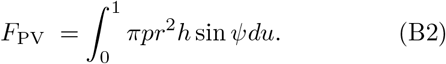

## Appendix C Elastic energy of the protein coat

Eq. (5) describes the elastic energy of the protein coat, which comes from the deviation of the normal curvature of the membrane surface *C*_*θ*_ in the *θ*-direction with the protein’s preferred curvature *C*_*p*_. It can be explicitly written as

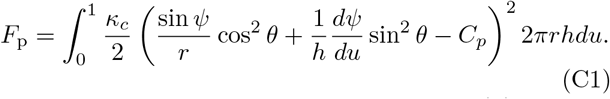

Since proteins only cover the region from *A*(*u*) = *A*_1_ to *A*(*u*) = *A*_2_ on the membrane, the bending rigidity of the protein coat *κ*_*c*_ as a function of *A*(*u*) takes the form

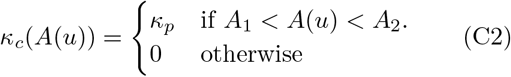

In practice, we approximate the piecewise function as

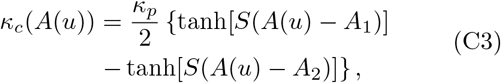

where *S* is a parameter to control the sharpness of the protein edge. Due to the dependence on *A*(*u*), it introduces inhomogeneity to the system.

**Appendix D: Variational equations of the composed system**

Combining the bending energy of the membrane (B1) and the proteins (C1), the pressure energy (B2), and the Lagrangian multipliers (A5), we arrive at the effective functional

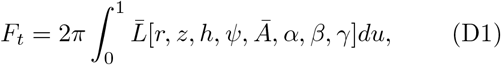

in which

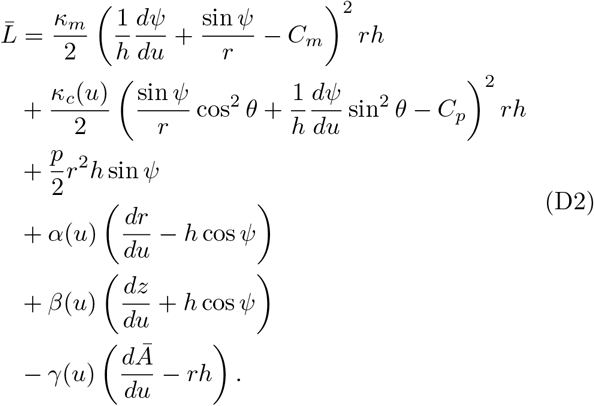

We perform variations against the functions {*ψ*(*u*), *r*(*u*), *z*(*u*), *Ā*(*u*), *α*(*u*), *β*(*u*), *γ*(*u*)} to obtain the Euler-Lagrange equations

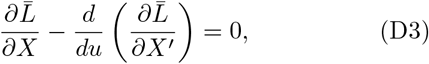

where *X* denotes any of the variable among *ψ*(*u*), *r*(*u*), *z*(*u*), *Ā*(*u*), *α*(*u*), *β*(*u*), *γ*(*u*) and *X*^′^ refers to their derivatives with respect to *u*. To be specific, major equations are

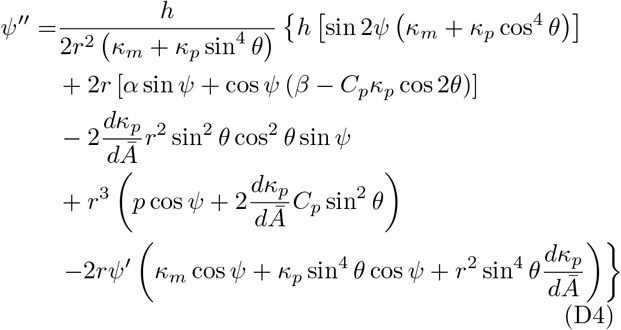

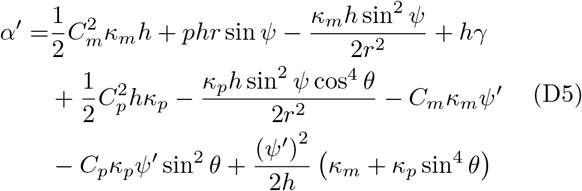

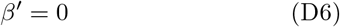

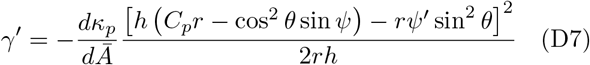

Note that the variation with respect to *h*(*u*) does not give an independent equation but a conserved quantity

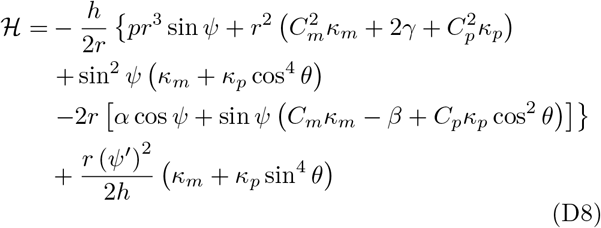

Meanwhile, we set boundary conditions as

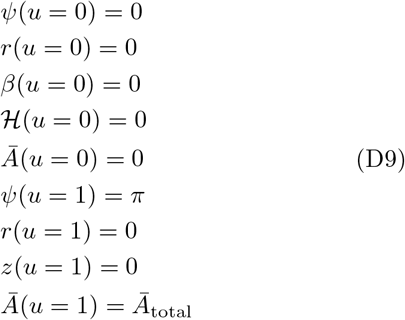

Eqs. (D4), (D5), (D6), (D7) and the geometric relations (A2), (A3), (A4) constitute the full set of equations we need to solve. They together make up 8 first order ordinary differential equations that define a boundary value problem with the unknown constant *h*, and the 9 boundary conditions given in Eq. (D9).

We solve the boundary value problem with the MAT-LAB solver bvp5c in an iterative manner. For instance, in order to calculate the shape transformations induced by varying the protein curvature *C*_*p*_, we start with *C*_*p*_ = 0 where the membrane is almost a sphere. Once we get the solution at *C*_*p*_ = 0, we can extend the solution to higher values of *C*_*p*_ with a number of small increment of Δ*C*_*p*_ and use the solution of the previous step as a guess for the current step. When there are multiple branches of solutions, continuously varying *C*_*p*_ could lead to error report due to no solutions beyond certain value of *C*_*p*_ (see the branch from A to B to C in Fig. 2b). When then change the iterative variable from *C*_*p*_ to *D*_*H*_ = *z*(0) *− z*(1) which is the pole-to-pole distance by adding a new BC in Eq. (D9). Accordingly, we set the protein curvature *C*_*p*_ as an unknown parameter. By alternating the iterative variables between *C*_*p*_ and *D*_*H*_, we can obtain the complicated free energy curve with Gibbs triangle.

## Notes

### Competing Interest Statement

The authors have declared no competing interest.

